# Noncannonical functions of Ku may underlie essentiality in human cells

**DOI:** 10.1101/2022.11.28.518141

**Authors:** Rachel D. Kelly, Gursimran Parmar, Laila Bayat, Matthew E.R. Maitland, Gilles A. Lajoie, David R. Edgell, Caroline Schild-Poulter

## Abstract

The Ku70/80 heterodimer is a key player in non-homologous end-joining DNA repair but has also been involved in other cellular functions like telomere regulation and maintenance, in which Ku’s role is not fully characterized. It was previously reported that knockout of Ku80 in a human cell line results in lethality, but the underlying cause of Ku essentiality in human cells has yet to be fully explored. Here, we established conditional Ku70 knockout cells to study the essentiality of Ku70 function. Endogenous Ku70 knockout was achieved using CRISPR/Cas9 editing in cells where Ku70 expression was maintained through integration of an HA-tagged Ku70 cDNA under the control of a doxycycline-inducible promoter. Ku70 conditional knockout cell lines were identified via western blotting, and edits were validated by Sanger sequencing. We visually observed cell death in Ku70 knockout cells 8-10 days post Ku70-HA depletion, and loss of viability following Ku depletion was quantified using crystal violet assays. Interestingly, assessment of telomere length in Ku70 knockout cells using telomere restriction fragment analyses did not reveal any changes in average telomere length following Ku70-HA depletion. Immunofluorescence analysis used to assess γH2AX foci accumulation as a measure of double-stranded DNA breaks following Ku70-HA depletion allowed us to conclude that increased DNA damage is not the driving cause of loss of cell viability. Finally, quantitative proteome analysis of Ku70 knockout cells following Ku70-HA depletion identified a number of pathways and proteins that are significantly dysregulated following the loss of Ku70, including processes which Ku function has been previously associated with such as cell cycle/mitosis, RNA related processes, and translation/ribosome biogenesis. Overall, this conditional Ku70 knockout system reveals that loss of Ku affects multiple cellular processes and pathways and suggests that Ku plays critical roles in other cellular processes beyond DNA repair and telomere maintenance to maintain cell viability.

**Author Summary:** The Ku70/80 heterodimer is a key player in non-homologous end-joining DNA repair, where it acts as a scaffold for other repair factors needed to process double-stranded DNA breaks. Ku has also been involved in other cellular functions like telomere regulation and maintenance, in which Ku’s role is not fully characterized. Previous data suggest that while loss of Ku70/80 can be tolerated in other species, Ku is essential to humans. We have established a conditional Ku70 knockout in HEK293 cells to evaluate the basis of Ku essentiality in human cells. While we observed loss of cell viability upon Ku depletion, we did not observe significant changes in telomere length nor did we record lethal levels of DNA damage upon loss of Ku, suggesting that the reasons for the loss of viability is not linked to the functions of Ku in DNA repair or at telomeres. Analysis of global proteome changes following Ku70 depletion revealed dysregulations of several cellular pathways including cell cycle/mitosis, RNA related processes, and translation/ribosome biogenesis. Our study reveals that loss of Ku affects multiple cellular processes and pathways and suggests that Ku plays critical roles in cellular processes beyond DNA repair and telomere maintenance to maintain cell viability.

## Introduction

One of the most hazardous forms of DNA damage that can arise from cellular processes are double-stranded DNA breaks (DSBs). Intracellular sources such as replication errors in dividing cells, reactive oxygen species formed as by-products of cellular metabolism, enzymatic action, and physical or mechanical stress can all lead to DSBs[1, 2]. Extracellular sources or environmental factors such as ionizing radiation, ultraviolet light, and chemical agents can also be a source for DSB formation. In mammalian cells, the primary method for repair of DSBs is the non-homologous end-joining (NHEJ) pathway, where repair factors work in synergy to directly ligate broken DNA[2].

Given the threat to genomic integrity that DSBs pose, efficient repair is a necessity for cellular survival. One of the first responders in the NHEJ pathway is a key protein known as Ku, which can arrive at the site of a break within seconds of the damage occurring[3, 4]. Ku is a heterodimer composed of two subunits, Ku70 and Ku80, and together the subunits form a ring-like structure that has high affinity for double-stranded DNA ends[5]. In the event of a DSB, Ku proteins will bind to each of the broken double-stranded ends in a sequence-independent manner[5, 6]. Once bound, the Ku heterodimer interacts with the DNA protein kinase catalytic subunit (DNA-PK_cs_) to form the DNA-PK complex that acts as a scaffold for other repair factors needed to ligate the DNA lesion[7]. Though the Ku heterodimer is best known for its role in NHEJ, it is also involved in other cellular processes. However, the precise functions of Ku in these pathways are not fully understood[8].

A few of Ku’s vital roles outside of NHEJ include V(D)J recombination and telomere regulation and maintenance[9, 10]. In mammals, single-stranded DNA overhangs at the ends of telomeres invade double-stranded repetitive telomeric TTAGGG sequences, and associate with six proteins of the shelterin complex to form structures known as t-loops[11, 12]. T-loops are essential to protecting DNA ends from being recognized as damage by DNA repair machinery, thus preventing chromosomal fusions and genomic instability[12].

Interestingly, loss of Ku appears to have different effects on telomere maintenance between species. In yeast, Ku binds to the RNA component of yeast telomerase (TLC1), specifically interacting with the stem loop of TLC1 to promote telomerase recruitment to telomeres, thus aiding in telomere lengthening[13]. Loss of Ku in yeast results in telomere shortening and can result in unwanted recombination between telomere ends[14]. In *Drosophila melanogaster*, a loss of Ku protein causes greater deprotection of telomere ends, leading to telomere lengthening that is observed in the absence of Ku[15]. In mammals, Ku has also been found to regulate telomere length. In mice, depletion of Ku results in both telomere lengthening and shortening, as well as increased chromosomal fusions[16–18]. Human Ku protein interacts with the telomerase RNA component (hTR) and the telomerase catalytic component (hTERT), and shelterin complex members[19, 20]. Human cells depleted of Ku display shortened telomeres and an increase in cell death[10,21,22]. A Ku80 knockout in human colon cancer HCT116 cells showed loss of telomere length, that was suggested to have occurred through formation of extrachromosomal circles of cleaved telomeric repeats known as t-circles[22, 23]. Telomere loss and cell death in HCT116 cells suggests that Ku may perform an essential role in human cells at telomeres[10, 22].

Homozygous knockout of either Ku70 or Ku80 subunits in mice causes a set of distinct phenotypic effects that are not displayed in heterozygous knockouts of Ku70/80[24, 25]. Characteristic phenotypes associated with Ku80 knockouts in mice include proportionally smaller body size, a loss of proliferating cells, longer cell doubling times, radiation sensitivity, deficiency in V(D)J rearrangement, and an arrest in the development of B and T lymphocytes[24]. Ku70 knockouts in mice resulted in similar deficiencies to Ku80 knockouts, but were also associated with a higher incidence of thymic tumours[25]. An interesting exception to the general observation that other species can tolerate a depletion of Ku protein is the fungus *Ustilago maydis.* A depletion of Ku in the fungus *U. maydis* has been shown to cause cell cycle arrest due to DNA damage response signaling at telomeres[26].

Although mice and other model organisms can tolerate loss of Ku and maintain viability, current evidence suggests that Ku knockout in human cells is lethal. Heterogeneous knockouts of Ku80 in HCT116 cells resulted in severe phenotypic effects, including defects in Ku DNA end-binding activity, sensitivity to ionizing radiation, and defects in cell proliferation, similar to the phenotypes of homozygous knockouts in mice[27]. Homozygous knockout of Ku80 in HCT116 cells resulted in loss of cell viability[22, 27]. A study using Nalm-6 cells did not report cell proliferation or telomeric defects following a heterozygous inactivation of either Ku subunit[28], but another study using the same cells found variability in the results previously reported[29]. The discrepancies between different cell lines and studies are not fully understood. Intriguingly, a more recent study reported that a deficiency of Ku protein led to an adaptive response where the Ku-deficient cancer cells exploited neighbouring cells to maintain their survival[30].

Given Ku’s ability to interact with shelterin complex members and telomerase components, as well as the severe telomeric shortening and loss of cell viability reported following the loss of Ku protein in human cells, it is possible that Ku is performing an essential function related to its action at telomeres. To investigate the function of Ku70, we created a conditional Ku70 knockout using CRISPR/Cas9 in TREx-293 cells. We find that loss of Ku70 protein levels directly led to a loss of cell viability, supporting the observation that Ku performs essential functions in human cells. Interestingly, decreased cell viability was not accompanied by critical loss of average telomere length, and did not appear to result from significant increases in unrepaired DSBs. Global quantitative proteomic analysis of whole cell extracts from Ku70 knockouts following depletion of Ku70 indicate that loss of Ku affects multiple cellular processes and pathways, and that Ku appears to play important roles beyond DNA repair and telomere maintenance in other cellular processes such as cell cycle and RNA-associated functions.

## RESULTS

### Generation of Ku70 Knockout Cells

To examine the impact of Ku70 knockout on cell viability, we first created a conditional TREx-293 cell line that expressed an inducible copy of the Ku70 cDNA. This was done to prevent the loss of cell viability if Ku70 was essential. TREx-293 cells were stably transfected with a doxycycline (Dox)-inducible exogenous copy of Ku70 cDNA using the Flp-In system (Flp-In™ T-Rex™, ThermoFisher). The exogenous copy of Ku70 was tagged at the C-terminus with a human influenza hemagglutinin (HA) sequence which allows monitoring of expression using an anti-HA antibody (exogenous Ku70 referred to as Ku70-HA henceforth). Following induction from the Tet-ON promoter with Dox, we tested the timeline of Ku70-HA depletion upon Dox removal (**Fig 1A**). Quantifications showed that, compared to Day 1, Ku70-HA protein abundance was significantly depleted by Days 4 (by ∼87%) while depletion reached ∼99% by Day 7 post Dox removal (**Fig 1B**).

**Fig. 1.**
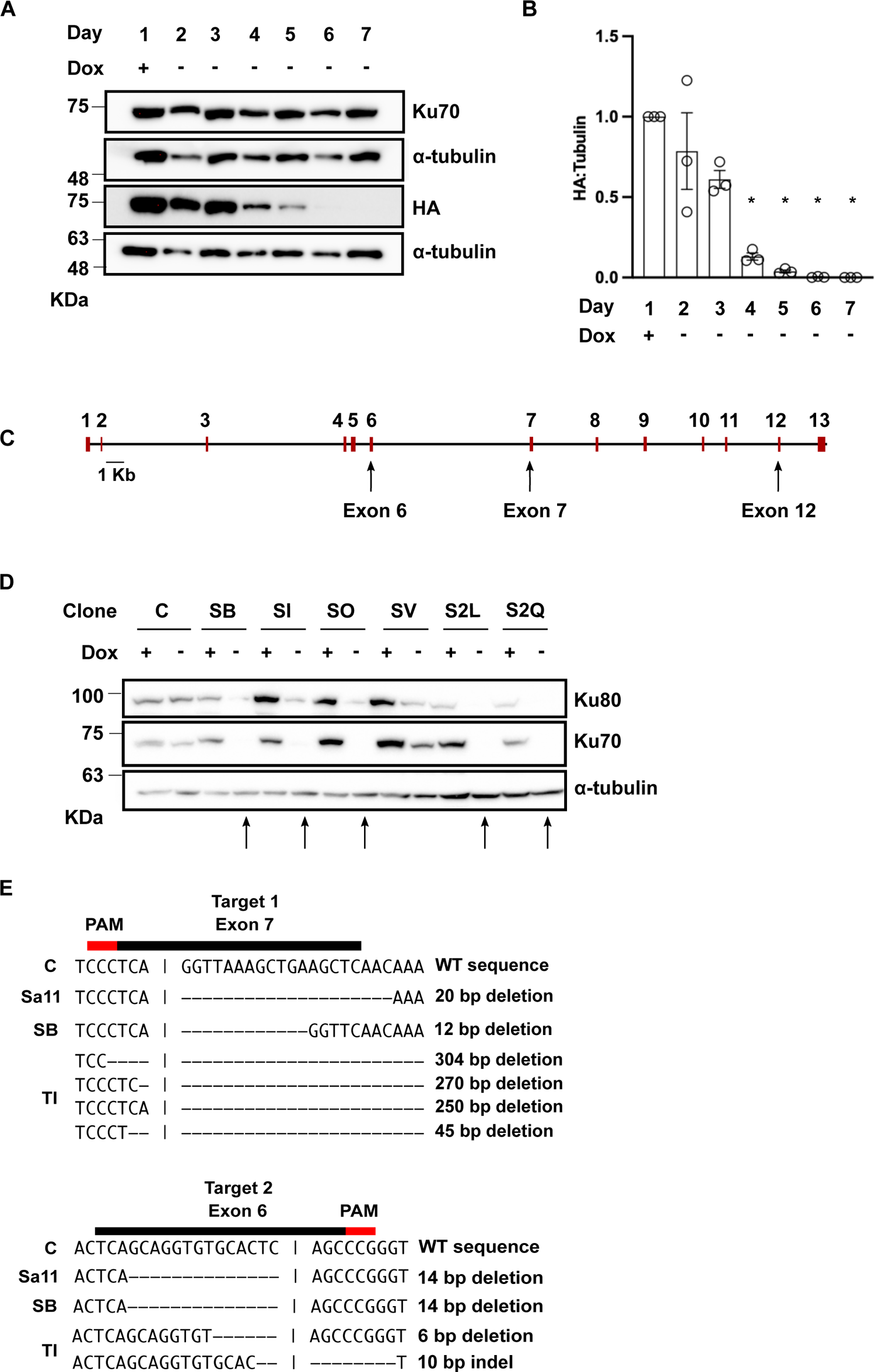
CRISPR Knockout Strategy, Screening, and Validation of Potential Ku70 Knockouts. **A.** Expression of Ku70 in TREx-293 cells after stable integration of exogenous Ku70-HA cDNA cells following Dox release. Extracts were collected from cells supplied with Dox (Day 1), and at subsequent days following Dox withdrawal (as indicated at the top). Extracts were run on SDS-PAGE and analyzed by western blot with the indicated antibodies. Both exogenous HA-Ku70 and endogenous Ku70 are detected by a Ku70 antibody and the depletion of Ku70-HA is tracked via HA tag. + indicates doxycycline (Dox) in cell media. - indicates Dox was removed from cell media. **B**. Quantification of the depletion of exogenous Ku70-HA normalized to alpha-tubulin Data are plotted as the mean of 3 biological replicates with error bars reporting+/- SEM. * indicates significant change compared to Day 1 Dox (p<0.005). **C.** Schematic of the CRISPR mediated knockout of endogenous Ku70 through targeting of exon/intron junctions at exons 7, 6, and 12 respectively. **D.** Analysis of Ku70 CRISPR knockout clones. Whole cell extracts from indicated clonal cells cultured in the presence (+) or absence (-) of Dox for at least 7 days were analyzed by western blot with the indicated antibodies. C indicates TREx-293 Ku70-HA control cells. Arrows indicate candidate knockouts. **E.** Mutations at target site 1 (Exon 7) and target 2 (Exon 6) for three Ku70 knockout clones. The wild-type sequence (C) is shown at top and dashes indicate indels found in the edited cell lines. The guide RNA is shown by the black bar above the sequence with the PAM sequence in red.

A CRISPR knockout strategy utilized three gRNAs simultaneously to target the exon/intron junctions of Ku70 exons 7, 6, and 12, respectively. The strategy of targeting the exon/intron junctions of the Ku70 gene was chosen to avoid off-target editing in Ku70 processed pseudogenes[8]. Targeting exon/intron junctions precluded Cas9 cleavage of Ku70 pseudogenes, or of the Ku70-HA. Editing was induced in the TREx-293 Ku70-HA cells with SaCas9 or the dual TevCas9 endonuclease[31] (**Fig 1C, S1 Fig**). After transfection with SaCas9 or TevCas9, colonies were screened by western blot for a reduction in endogenous Ku70 after Day 7 post Dox withdrawal (**Fig 1D**). From this screening, 27 potential Ku70 knockout clonal cell lines were identified. Of the 27 Ku70 knockouts, 18 were edited with Cas9 endonuclease cleavage, and 9 were edited with TevCas9. Edits at the three target sites were validated by T7 endonuclease assays and Sanger sequencing of PCR products encompassing the editing sites (**Fig 1E**, S1 Data). Three clonal cell lines (Sa11, SB, and TI) that had insertions or deletions at two target sites were chosen for further characterization (**Fig 1E**).

### Ku70 knockout cells lose viability 8-10 days post exogenous Ku70-HA withdrawal

To establish a timeline for viability in Ku70 knockout cells as Ku70-HA is depleted, Dox release curves were generated to determine the amount of time between reduction in Ku70-HA protein and cell death for Ku70 knockout clones. We examined Ku70-HA protein levels by western blot in one knockout cell line (SB), finding that Ku70-HA depleted to ∼1% of the Day 1 amount by Days 6 and 7 post Dox withdrawal (**Fig 2A and 2B**). Viable Ku70 knockout cells decreased between 8-10 days post Dox withdrawal. By Day 8 post Dox withdrawal, cells displayed a condensed, rounded phenotype, and ∼60% of the cells had begun to lift off the plate as compared to Day 1 (**Fig 2C**).

**Fig. 2.**
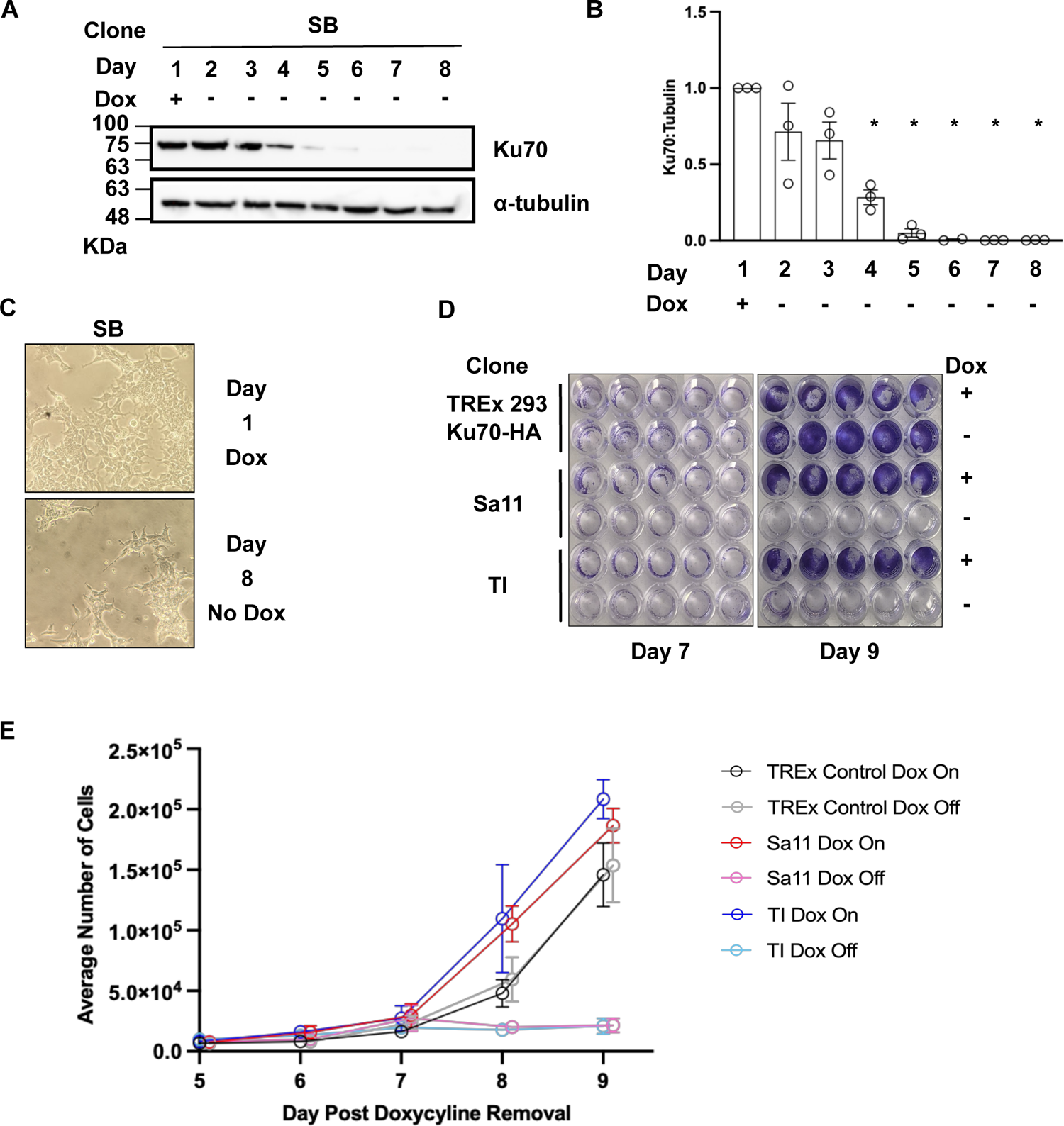
Loss of cell viability following depletion of Ku70 in conditional Ku70 knockout clones. **A.** Western blot of Dox depletion curve for the SB Ku70 knockout clone on the indicated days. Whole cell extracts from SB cells cultured in presence (+) or absence (-) of Dox and were analyzed by western blot with the indicated antibodies. **B.** Quantification of Ku70 relative to alpha-tubulin plotted as the mean of 3 biological replicates with error bars reporting+/- SEM. * indicates Ku70 is significantly changed compared to Day 1 Dox (p<0.05). **C.** Cell morphology of Ku70 knockout cells maintained in Dox and on Day 8 post Dox withdrawal. Cells were visualized by phase-contrast with a 20X magnification. **C.** Images of cells stained with crystal violet fixed at Day 7 and Day 9 post Dox withdrawal. **D.** Crystal violet assay assessing cell viability following loss of Ku70 expression. TREx-293 Ku70-HA Control cells, and two Ku70 knockout clones, Sa11 and TI were cultured with Dox (Dox On) or without Dox (Dox Off) and plated on 96-well plate at Day 5. Cells were fixed and stained at days 5 to 9. **E.** Crystal violet assays were quantified and plotted (n=3 for each time point). All points are nudged 0.1 along x-axis to allow differentiation between samples.

Crystal violet assays were used to quantify loss of cell viability post Ku70-HA withdrawal. We plated the TREx-Ku70-HA Control cells, and the Sa11 and TI Ku70 knockout cells on Day 5 post Dox withdrawal along with growth-matched controls of each of the three cell lines maintained in Dox-containing media. Cells were fixed at 24-hour timepoints starting at Day 5 post Ku70-HA withdrawal (0h) and ending at Day 9 post Ku70-HA withdrawal (96h). Ku70 knockout cells grown without Dox displayed a significant reduction in the number of viable cells adhered to the wells of the plate as compared to growth-matched controls (**Fig 2D**). For the Sa11 knockout clone, by Day 9 post Ku70-HA Dox withdrawal, only ∼11.6% of the average number of cells were still adhered compared to the Sa11 Dox On control sample. Similarly, for the TI knockout, only about ∼10% of cells remained 9 days after Dox removal compared to the TI Dox On control (**Fig 2E**). Collectively, this data indicated that the loss of Ku70 correlated with a severe decrease in cell viability.

### Ku70 knockout cells do not undergo significant changes in telomere length following exogenous Ku70-HA withdrawal

Previous studies showed that in HCT116, HeLa, and Nalm-6 cells, loss of Ku protein resulted in telomere shortening[29,32,33]. We therefore investigated the telomere status of cells in which Ku70 was depleted. We chose to evaluate average telomere length at Day 8 post Dox withdrawal because Ku70-HA was maximally depleted and cells began to lose viability. Average telomere lengths of TREx-293 Sa11, SB, and TI Ku70 knockout clones were assessed using a telomere restriction fragment analysis (**Fig 3A**). In the SB Ku70 knockout clone, there was an average telomere length of 3.1 Kb on Day 1 +Dox and 4.3 Kb on Day 8 no Dox (p=0.9274). For another Ku70 knockout clone, Sa11, there were also no significant changes in telomere length identified (3.9 Kb on Day 1 Dox and an average length of 4.5 Kb on Day 8 Dox withdrawal (p=0.9998). For the final clone analyzed TI, the average telomere length on Day 1 Dox was 4.0 Kb and on Day 8 no Dox was 3.6 Kb (p>0.9999). The difference in telomere lengths for Ku70 knockouts were also not significantly different from control cells that did not undergo Ku70 knockout (TREx-293 Ku70-HA) either at Day 1 (3.3 Kb), Day 8 (3.6 Kb), and No Dox Day 8 (3.6 Kb). There was also no significant difference in average length compared to TREx-293 cells lacking the exogenous Ku70-HA exogenous vector (4.6 Kb). Overall, the data show that by Day 8, when Ku expression is diminished and cell viability starts to be compromised, there is no significant change in average telomere length when compared to Day 1 when Ku levels are unaffected (**Fig 3B**). This data suggests that Ku70 depletion is not associated with telomere shortening in HEK293 cells.

**Fig. 3.**
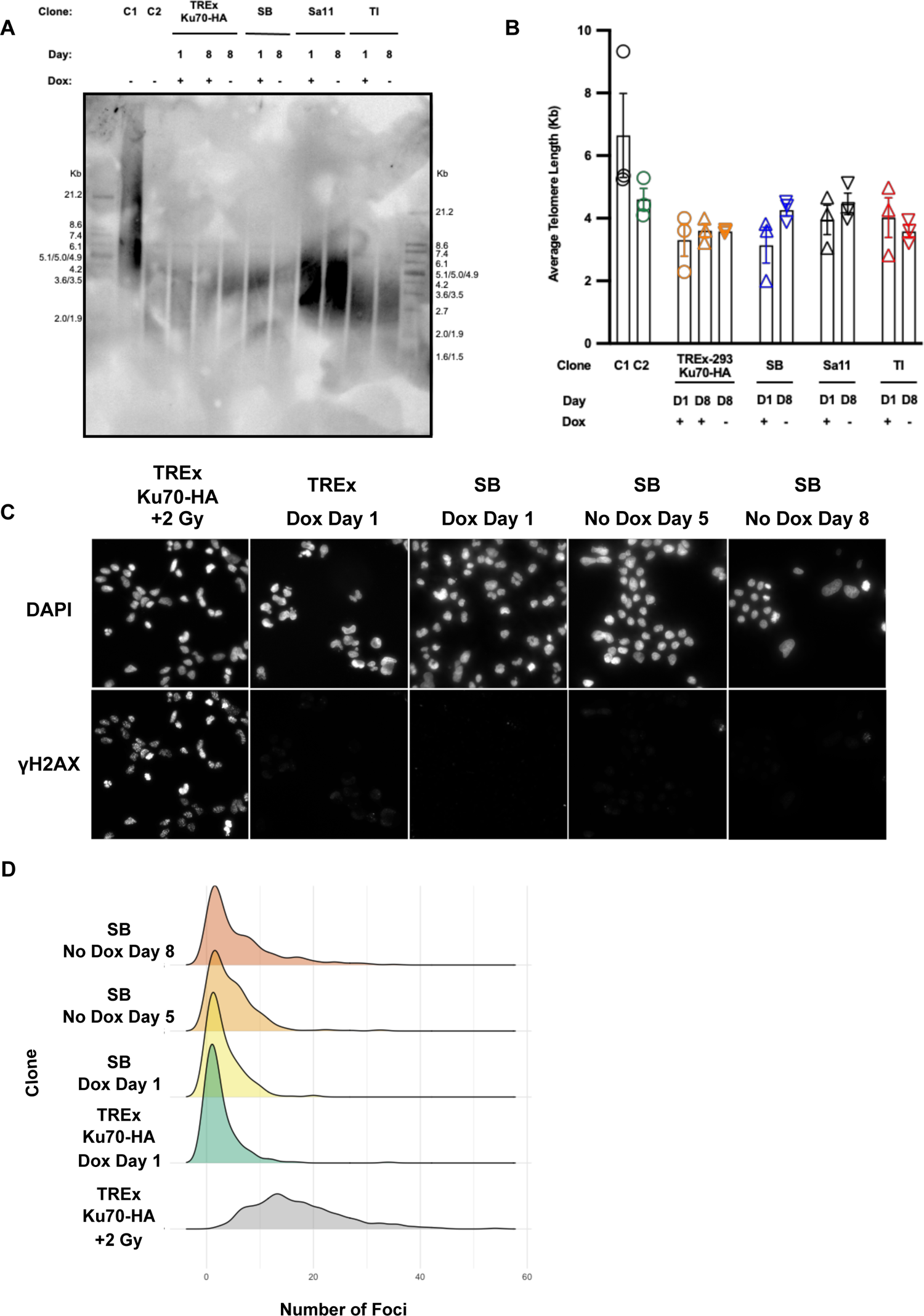
Cell viability in Ku70 knockouts is not correlated with telomere shortening or γH2AX foci accumulation. **A.** Representative telomere restriction fragment (TRF) analysis of control cells compared to Ku70 knockout cells (clones SB, Sa11, TI) following exogenous Ku70 depletion on the days indicated. C1 denotes TRF kit control DNA (U937 cells). C2 denotes unedited TREx-293 cells. TREx Ku70-HA denotes unedited TREx-293 cells with Dox-inducible exogenous Ku70. +/- indicate the presence or absence of Dox in cell media. Size markers are indicated on the side. **B.** Average telomere length measurements from TRF analyses (N=3). Day indicates what day samples were collected during Dox depletion curve. + or - indicates presence or absence of Dox in cell media. C1 is a control cancer cell line. C2 is TREx-293 cell line. **C.** Immunofluorescence images displaying γH2AX foci in Ku70 knockout cells on the days indicated in media containing Dox or following Dox withdrawal (no Dox). TREx-293 Ku70-HA cells treated with 2 Gy of ionizing radiation act as a positive control. **D.** Density plots representing average number of foci per cell nucleus for each condition/treatment analyzed (N=3).

### Examination of γH2AX repair foci accumulation in Ku70 knockout cells

We considered the possibility that loss of cell viability could be due to an accumulation of unrepaired DSBs in the Ku70 knockout cells. Immunofluorescence was used to examine γH2AX foci accumulation, a marker of DSBs, following depletion of Ku70. Previous work demonstrated that in the absence of Ku80 protein, there is a significant increase γH2AX foci, a marker of DSBs, in knockout cells[34]. We analyzed SB Ku70 knockout cells on Day 1 Dox, and Days 5 and 8 post Dox removal and compared it to TREx-293 Ku70-HA control cells that were treated with 2 Gy of ionizing radiation (IR) (**Fig 3C**). The number of foci per nucleus increased from an average of 3.1 foci/nucleus on Day 1 to 7.2 on Day 8 No Dox (S2 Fig). The average number of foci/nucleus for SB Ku70 knockout cells on Days 1 Dox, and Days 5 and 8 post Dox removal were found to be significantly lower compared to TREx-293 Ku70-HA control cells that were treated with 2 Gy of ionizing radiation (18.7 foci/nucleus). However, no significant differences were found between SB Ku70 knockout cells at the different days analyzed post Ku70-HA depletion. Density plots of the number of foci/nucleus show a higher number of cells with more γH2AX foci in SB Ku70 knockout cells by Day 5 and Day 8 post Ku70-HA removal compared to Day 1 (**Fig 3D**). Despite this general trend, the majority of nuclei post Ku70-HA depletion contain low numbers of γH2AX foci that are similar to cells maintained in Dox and unedited TREx-293 Ku70-HA cells. Overall, our data show that the elevated amount of γH2AX foci observed in absence of Ku does not reach the level induced by 2 Gy of IR which was reported to result in more than 70% survival using a colony forming assay in HEK293 cells[35]. These findings lead us to conclude that it is not the accumulation of DSBs that is the driving factor behind the loss of cell viability in Ku70 knockout cells.

### Proteomic analysis of global protein abundance changes in Ku70 knockout cells

To identify pathways that are affected by the loss of Ku expression, we sought to evaluate the proteomic changes that occur upon Ku depletion. Whole cell extracts of SB cells subjected to Dox withdrawal and the growth-match controls cultured with Dox on Days 1, 4, 6 and 7 (N=3) were selected for proteome analysis by mass spectrometry. These days were chosen because by Day 4 post Dox withdrawal, the relative amount of Ku70 was reduced significantly (∼30% of the amount on Day 1), and the relative amount of Ku70 was ∼1% of the Day 1 amount by Days 6 and 7 post Dox withdrawal, which occurs before cells are lifting from plates on Day 8 post Dox removal.

Using label-free quantification, 5353 proteins were quantified in at least 3 samples (S2 Data). In agreement with the western blot data, Ku70 protein abundance gradually decreased after Dox removal (**Fig 4A**). By Day 4 following Dox withdrawal, Ku70 (XRCC6) protein levels were decreased to approximately half of those recorded at Day 1 and were also significantly decreased compared to the Day 4 +Dox growth-matched control. This trend continued to widen to ∼0.69 and ∼0.97 mean differences for Day 6 and Day 7 Ku-depleted to +Dox growth-matched controls, respectively (**Fig 4A**). A similar trend was observed for Ku80 relative protein abundance with mean differences of 0.34 between Dox-depleted and control Day 4 samples, 0.69 on Day 6, and ∼1 on Day 7 (**Fig 4A**).

**Fig. 4.**
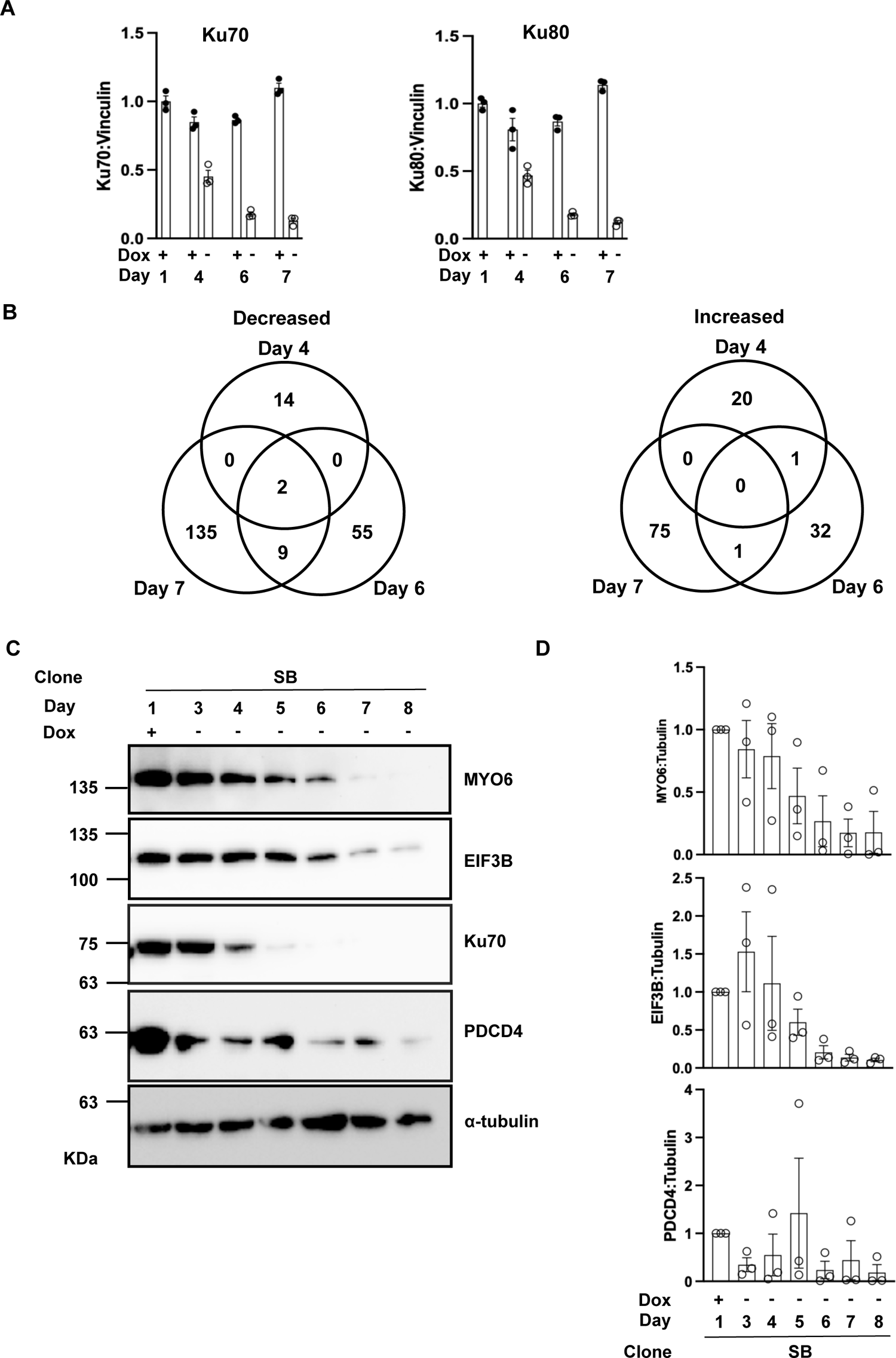
Proteomic analysis following depletion of Ku and validation of altered proteins. **A.** Quantification of relative abundance of Ku70:Vinculin protein LFQ intensities in SB clone samples from cells maintained in Dox (+) or without Dox (-) at the indicated days in culture. Ku70 relative abundance was set at 1 at Day 1. +indicates Dox is added to cell media. - indicates Dox has been removed from cell media. **B.** Quantification of relative abundance of Ku80:Vinculin protein LFQ intensities compared to the Day 1 control, as plotted in A. **C.** Venn diagram of the number of proteins found to be decreased or increased significantly on Days 4, 6, and 7 post Dox withdrawal (Fold-change ≥ 1.5; p-value ≤ 0.05). **D.** Western blot validation of proteomic analysis results for 3 candidate proteins at the days listed post Dox withdrawal for the SB Ku70 knockout clone. **E.** Quantification of MYO6, EIF3B, and PDCD4 protein levels relative to alpha-tubulin for (N=3) western blots. +/- indicates the presence or absence of Dox from cell media. Day post Dox withdrawal are indicated.

We next examined global proteome changes, focusing on Day 4, Day 6, and Day 7 comparisons. The only proteins depleted on all days examined were Ku70 and Ku80. On Day 4 post Dox withdrawal, 21 proteins were significantly increased ≥1.5 fold-change (FC) compared to the growth-matched controls and 16 proteins were decreased ≥1.5 FC (**Fig 4B**; S3 Data, S3 Fig). By Day 6, 34 proteins were significantly increased ≥1.5 FC and 66 proteins were decreased ≥1.5 FC (**Fig 4B**). On Day 7 post Dox withdrawal, there were 76 proteins increased ≥1.5 FC and 146 proteins were decreased ≥1.5 FC (**Fig 4B,** S3 Data). A student’s two-way t-test determined that 12 proteins were significantly altered with a Q value of 0.05 or less on Day 7 (**Table 1,** S4 Data).

**Table 1.** Proteins Significantly Changed

| **Protein names** | **Gene names** |
| --- | --- |
| X-ray repair cross-complementing protein 6 | XRCC6 |
| X-ray repair cross-complementing protein 5 | XRCC5 |
| Eukaryotic translation initiation factor 5 | EIF5 |
| Eukaryotic translation initiation factor 3 subunit B | EIF3B |
| Bystin | BYSL |
| Little elongation complex subunit 2 | ICE2 |
| S1 RNA-binding domain-containing protein 1 | SRBD1 |
| ADP-ribosylation factor GTPase-activating protein 1 | ARFGAP1 |
| Ubiquitin carboxyl-terminal hydrolase 33 | USP33 |
| Phosphatidylinositol glycan anchor biosynthesis class U protein | PIGU |
| Stromal interaction molecule 2 | STIM2 |
| 2-5-oligoadenylate synthase 3 | OAS3 |

Three candidate proteins, MYO6, PDCD4, and EIF3B, were chosen to validate the quantitative proteomic results based on if they were significantly altered (either increased or decreased) on Day 6 and Day 7 with a ≥1.5 fold-change and p-value ≤0.05. Western blots for relative protein abundance confirmed that there is a general trend of decreasing protein abundance for MYO6, PDCD4, and EIF3B for the SB Ku70 knockout clone (**Fig 4C**). Quantification of the three candidate proteins in relation to alpha-tubulin showed that the mean protein abundance had depleted to ∼6.8% of the Day 1 Dox abundance for EIF3B by Day 8 (**Fig 4D**). Similarly, by Day 8 post Dox removal MYO6 showed a decrease in protein abundance of ∼1.5% the mean of Day 1, and ∼1.7% of the mean relative abundance for PDCD4 (**Fig 4D**). These results were also validated by western blotting with extracts from another Ku70 knockout clone, Sa11 (S4 Fig). Overall, these data provide validation of proteomic analysis results.

Next, we evaluated pathways affected by the loss of Ku expression using Metascape. From the lists of proteins significantly altered (FC ≥1.5, p-value ≤ 0.05), the top 10 biological pathway networks were visualized for each day of analysis (Day 4, Day 6, and Day 7) using a heatmap (**Fig 5,** S5 Data). Enriched biological terms associated with decreased proteins featured on the heatmap were coloured according to p-value. Some of the networks associated with decreased protein abundances have been previously associated with Ku function including apoptosis, and other pathways involving mitosis or the cell cycle have been implicated with Ku, although the precise function of Ku in these cellular processes is not yet fully understood. Interestingly, the analysis identified pathway networks in which Ku’s function is not well established, such as metabolism of lipids (**Fig 5A**). Notable networks with upregulated proteins included ncRNA metabolic process, and cell cycle G2/M transition phase, which have been previously implicated in Ku function[8, 26] (**Fig 5B**).

**Fig. 5.**
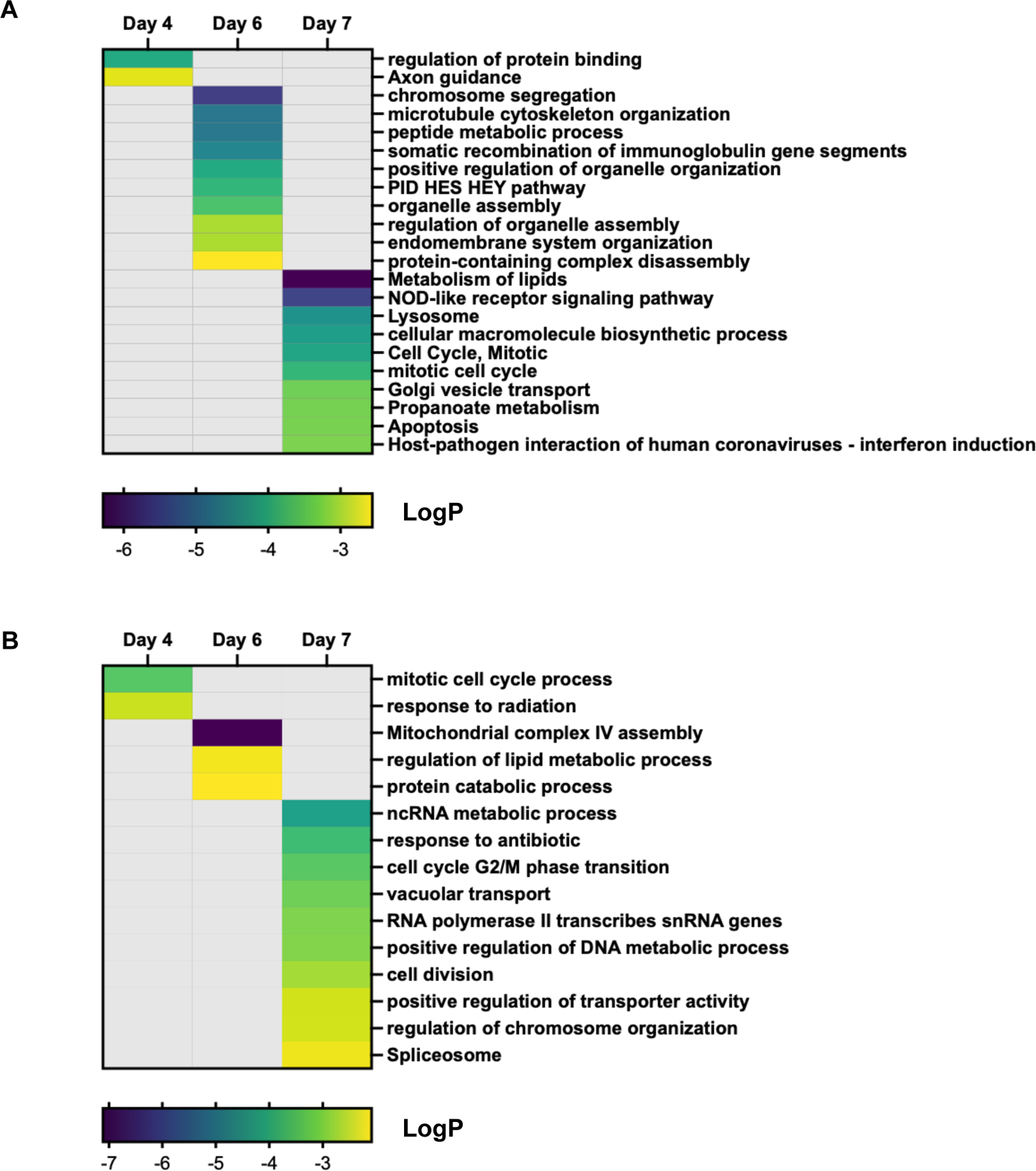
Global Proteomic Changes in Ku70 Knockout Cells May Indicate Non-Canonical Essential Function for Ku70. **A.** Enriched biological terms associated with significantly decreased proteins from proteomic analysis on days 4, 6, and 7 no Dox (FC ≥ 1.5, p-value ≤0.05). **B.** Enriched biological terms associated with significantly increased proteins from proteomic analysis on days 4, 6, and 7 no Dox (FC ≥ 1.5, p-value ≤0.05). Pathway and process enrichment analysis was performed via Metascape and the top 10 enriched terms for each day are displayed in a heatmap (p-value < 0.01, minimum count of 3, enrichment factor > 1.5). Enriched terms are coloured according to p-value. The p-values are displayed as LogP.

## Discussion

We created a conditional Ku70 knockout system using TREx-293 cells to investigate Ku’s essentiality in human cells. In this system, we expressed a Dox-inducible exogenous Ku70-HA and subsequently knocked out endogenous Ku70, allowing precise monitoring of Ku70 depletion upon Dox removal. We determined that loss of Ku70 (and its obligate heterodimer partner Ku80) resulted in cell lethality that occurred shortly after Ku depletion. Cells were nonviable and lifted from plates 8-10 days post removal of doxycycline from the media, at which time the relative amount of Ku70-HA was reduced to 1% of the initial level prior to Dox removal. The average telomere length of each of the knockout clones did not change significantly upon Ku70-HA depletion, and was not significantly different from the average telomere lengths of unedited cells. Analysis of global proteomic changes between control (Dox on) and Ku-depleted (Dox off) for the SB Ku70 knockout clone do not show major pathway changes or protein abundance changes in relation to telomere regulation or maintenance, but other vital cellular processes are impacted including cell cycle and RNA metabolism.

Of the 27 Ku70 knockout clones established, 18 were edited using SaCas9 and 9 were edited with the TevCas9 fusion endonuclease. It has been previously established that SaCas9 and TevCas9 editing events can result in large deletions of genomic sequences that are over 2 Kb in length[36], and it is possible that the use of multiple gRNAs in conjunction with these endonucleases contributed to the heterogenous types of edits identified, including larger indels. It was also of note that the gRNA for Target 3 in Exon 12 was inefficient for editing and didn’t contribute to the creation of Ku70 knockouts characterized in this study. It is also possible that editing by Cas9 at Target 3 in exon 12 does not produce knockouts, and that is why indels were not identified at this exon/intron junction in the knockout clones screened.

Previous studies found that the knockout or knockdown of Ku protein resulted in dramatic telomere shortening in telomerase positive cells[29,32,33,37], but this is not consistent with our findings in TREx-293 cells, which are also telomerase positive. A dramatic loss of telomeric TTAGGG repeats could cause a critical telomere length to be reached where cell cycle arrest and apoptosis is initiated[38], but we did not find a significant difference in telomeric length compared to unedited cells. Our results parallel those of Uegaki et al. who reported that a heterozygous inactivation of Ku70 or Ku80 in telomerase-positive Nalm-6 cells did not result in significant telomere shortening[28]. Experiments involving Ku70/80 knockdown in human cells that do not rely on telomerase, but another method of telomere length regulation known as alternative lengthening of telomeres (ALT), also did not display changes in average telomere length following knockdown of Ku[39]. In accordance with findings from previous literature[21,29,37,39], the Ku70 knockout human cells we have generated lose viability following depletion of the exogenous Ku70-HA protein. Collectively, these data suggest that the dramatic loss of telomere length seen in previous studies may be a specific phenotype due to variations between cell lines, and that the essential function at Ku may not be due to telomere length regulation. Moreover, since a critically low telomere length can induce a DNA damage response (DDR), we would have expected to observe significant induction of DDR proteins such as p16 or p53, but our proteomic analyses did not show significant changes in expression that would indicate a response activated by dysfunctional telomeres.

Expectedly, loss of Ku induced an increase in γH2AX foci as previously reported[37], due to the loss of NHEJ repair of basal levels of DSBs. However, the elevated amount of γH2AX foci observed in absence of Ku did not reach the level induced by 2 Gy of IR which has been reported to result in about 30% cell death as assessed by colony formation ability in HEK293 cells[35]. These findings lead us to conclude that it is not the accumulation of DSBs that is the driving factor behind the loss of cell viability in Ku70 knockout cells.

In examining global proteomic changes following Ku70 depletion, we first noted that the largest number of protein changes greater than 1.5 fold-change was seen on Day 7 post Ku70-HA withdrawal compared to control Day 7 samples. This finding is in line with the observation that Day 7 had the largest mean difference in relative abundance of both Ku70 and Ku80. Almost double the number of proteins were down regulated 1.5 FC or greater on Day 7 post Dox withdrawal compared to the number of proteins upregulated. This difference may be due to the onset in cell death occurring, as the degradation of cellular components is a key step in apoptosis[40]. One of the proteins upregulated on Day 7 in Ku70 knockout cells is MTCH1 (Mitochondrial Carrier 1), also known as PSAP (presenilin 1-associated protein), a mitochondrial protein that has been shown to induce apoptosis when overexpressed in HEK293 cells[41], and has been more recently shown to have two isoforms that are proapoptotic[42]. Downregulation of another protein found on Day 7, TIGAR, was also shown to induce cell death through accumulation of reactive oxygen species[43, 44]. The combined observations, along with the reduction in anti-apoptosis Bcl-2 family member MCL1[45] protein levels on Day 6, provide evidence that these cells undergo apoptosis as Ku70 protein levels deplete in the conditional knockout cells. Interestingly, MCL1 has also been identified as an inhibitor of the Ku complex, capable of inhibiting NHEJ DNA repair to facilitate homologous recombination[45].

Of the 10 proteins found dysregulated on Day 7 post-Dox removal, two of them, eukaryotic translation initiation factor 5 (EIF5) and eukaryotic translation initiation factor 3 subunit B (EIF3B), are initiation factors for the translation of proteins. Translation initiation and proper formation of the 80S ribosomal initiation complex depends on GTP hydrolysis by EIF5 and coordinated action of other translation initiation factors like EIF3B[46]. More recent studies have also shown that knockdown of EIF3B can inhibit cell cycle progression and proliferation in cancer cells[47, 48]. Interestingly, two other proteins from the list of significantly dysregulated proteins are also involved in translation. Bystin is a protein that works to promote cell proliferation through formation of the 40S ribosomal subunit[49], and S1 RNA binding domain 1 (SRBD1) containing proteins are also predicted to be involved in ribosome biogenesis[50]. It is interesting that several factors associated with translation and ribosome biogenesis were significantly dysregulated compared to control cells in our study, as Ku has been previously implicated with RNA binding[8] and more specifically, it has been implicated as an interactome member of RNA Polymerase I and RNA involved in ribosome biogenesis[51]. Ku has also been implicated in rRNA processing via the DNA-PK complex[52].

The functions of the other top proteins found to have significantly altered protein levels include STIM2 (Stromal Interaction Molecule 2), which is associated with calcium release in the endoplasmic reticulum that can have an effect on multiple cellular processes[53]. OAS3 (2’-5’-Oligoadenylate Synthetase 3) acts to restrict viral replication and OAS family members can act as a dsRNA sensor[54]. Ku has also been previously shown to interact with hairpin structure of RNA[55], and given the relatively new field of Ku research and RNA biology, it is therefore possible Ku might interact directly or indirectly with other proteins participating in this sensing system directly or indirectly. ADP ribosylation factor 1 GTPase activating protein 1 (ARFGAP1) is involved in membrane trafficking[56] and the relevant biological terms that were found in our Metascape analysis also reflect vesicle transport as a pathway being dysregulated. Mitotic cell cycle checkpoints were found to be significantly changed in our pathway analysis, and from the list of proteins significantly changed between controls and experimental samples, USP33 (Ubiquitin Specific Peptidase 33) is involved in mitosis and cell division control[57]. Ku has been previously implicated to play a role in the G1/S and G2/M checkpoint phases of mitosis, as downregulation of Ku in *U. maydis* resulted in cell arrest at the G2/M checkpoint[26], and another study noted G2/M defects in Ku-deficient hamster cells treated with a a DNA topoisomerase II inhibitor[58]. Also, reduction of Ku80 protein levels in human cells was reported to trigger an accumulation of cells halted at the G1/S transition[59].

MYO6 is a motor protein implicated in intracellular vesicle and organelle transport, and the depletion of this protein has also been shown to affect cell proliferation/cell cycle progression and result in increased apoptosis in colon cancer cells[60] and prostate cancer cell lines[61]. The MYO6 interactome identified through BioID experiments is linked to multiple cellular processes, including centrosomal proteins that operate in organizing microtubules and have key roles during mitosis[62]. One of the BioID interaction network members with MYO6 is PCM1[62], which is also a significantly dysregulated protein identified in our proteomic analysis results that is associated with centrosomal functions. PDCD4 is an inhibitor of apoptosis, and has been shown to increase cell sensitivity to apoptosis[63]. The decrease in EIF3B, MYO6, and PDCD4 observed in the proteomic analysis and in our validations using western blots could be contributing to the loss of cell viability and dysregulation of cell cycle that is noted in the proteomic analyses.

Taken together, our results support that the Ku heterodimer does play an essential role in human cells and maintaining cell viability. Interestingly, our results indicate that Ku’s essential role in humans is not exclusively due to its action in maintaining telomere length. Our global proteomic analysis showed that a number of essential cellular processes, such as ribosome biogenesis/translation, RNA interactions, and mitotic cell cycle control are dysregulated in the absence of Ku. The conditional Ku70 knockout system developed here will allow us to evaluate more precisely the molecular links between Ku70 and the identified proteins, and how these relationships contribute to Ku essentiality in human cells.

## MATERIALS AND METHODS

### Plasmid Constructs

px458SpCas9_GFP_ (SpCas9-2A-GFP) and px459SpCas9_PuroR_ (SpCas9-2A-puro) vectors were previously obtained from Feng Zhang through Addgene (Addgene plasmid # 48138 and plasmid # 62988) for transfection into mammalian cell lines[64]. The nuclease, SpCas9, is linked to green fluorescence protein (GFP) and a puromycin resistance marker, respectively. The SaCas9 construct was created by cloning the full length SaCas9 into px458SpCas9_GFP_ following excision of the SpCas9 insert. Polymerase chain reaction (PCR) amplification of the pac gene encoding puromycin N-acetyl-transferase from px459SpCas9_PuroR_ was used to clone puromycin resistance into this construct to create px458_PuroR_SaCas9_GFP_. px458_PuroR_TevSaCas9_GFP_ was constructed by cloning I-TevI (amino acids 1–169) in front of the N terminus of SaCas9. The pBIG2R-Ku70 tetracycline repressible plasmid was created by cloning full length Ku70 into the multiple cloning site of the pBIG2r vector[65]. An HA-tag was subcloned to the C-terminus of Ku70 in pBIG2R-Ku70. To create pcDNA5/FRT/TO-Ku70-HA, for the TREX Ku70-HA tetracycline repressible system, full length Ku70-HA from pBIG2R-Ku70-HA was PCR-amplified using primers containing restriction enzyme sites and cloned into pcDNA5/FRT/TO.

### Designing gRNA

Since Ku70 has five pseudogenes that contain coding sequences from endogenous Ku70, gRNA was designed to target intron-exon junctions in Ku70. A script was used to locate potential Cas9 and TevCas9 target sites in Ku70. This script searched the Ku70 DNA sequence for regions that spanned intron-exon junctions and had the consensus sequence required for Tev nuclease and Cas9 nuclease cleavage. The consensus sequence for SaCas9 and TevSaCas9 target sites was 5‘ CNNNG(N)_34-40_NNGRRT 3’. 5‘ CNNNG 3’ is the consensus sequence required for Tev nuclease cleavage. 5‘ NNGRRT 3’ is the PAM sequence for SaCas9 required for Cas9 cleavage.

Ku70 knockout was carried out using following gRNAs:

SaCas9 & TevSaCas9

Target 1: 5’ AGCTTCAGCTTTAACCTGA 3’

Target 2: 5’ ACTCAGCAGGTGTGCACTCAGC 3’

Target 3: 5’ TCATTGCTTCAACCTTGGGCAC 3’

All of the target sites chosen spanned both intronic and exonic region of the Ku70 gene. TevCas9 had an additional cut site present upstream of the gRNA in these target sites determined by the Tev nuclease consensus sequence of 5’ CNNNG 3’.

gRNAs were ordered in the form of synthesized oligonucleotides with *Bbs*I cut site compatible overhangs added to each side. The designed gRNA was cloned into px458SpCas9_GFP_, px458TevSpCas9_GFP_, px459SpCas9_PuroR_, and px459SpCas9_PuroR._ This was accomplished using Golden Gate assembly, following the protocol outlined in Engler *et al.*, (2008)[66]. However, the restriction enzyme *Bbs*I was used instead of *Bsa*I. After Golden Gate assembly, heat shock transformation was performed using *Escherichia coli* (DH5α) Plasmids were purified using EZ-10 Spin Column Plasmid DNA Miniprep Kit by (Bio Basic Inc). Correct gRNA insertion was confirmed by DNA sequencing.

### Cell Culture, treatments, and transfections

HEK293 TREx cells (Invitrogen Canada Inc.) were cultured in high-glucose Dulbecco’s modified Eagle’s medium (DMEM) supplemented with 10% fetal bovine serum (FBS) at 37 °C in 5% CO_2._ to which 1% L-glutamine, and 1% sodium pyruvate were added. Transfections were performed using jetPRIME Versatile DNA/siRNA transfection reagent, following the manufacturer’s instructions (Polyplus Transfection Inc). Antibiotic was added 24-48 hours after transfection for selection. Single clones that grew in the presence of antibiotic were moved to 96 well plates and then grown until they could be moved to 6-well plates. Clones were assessed by western blot for a reduction in Ku70 protein following Dox withdrawal for at least 7 days.

Ku70^-/-^ cell lines were maintained with 1μg/mL Doxycycline (BioShop Canada Inc.) administered every 48 hours, and 15 μg/mL Blasticidin (MULTICELL), and 15 μg/mL Hygromycin (MULTICELL) which were administered every 96 hours.

### Exogenous Ku70 Depletion Curves and Western Blotting

In 6-well tissue culture dishes, 300,000 cells were plated per well. Ku70^-/-^ cell lines were supplemented with 1μg/mL Doxycycline (BioShop Canada Inc.) in cell media the day before the cells were plated onto the 6-well tissue culture dishes without Doxycycline. Cells were split on Day 3 and Day 5 1:3. Cells were trypsinized, collected, and the pellet was washed with phosphate-buffered saline (PBS; Wisent). Whole cell extract of cell pellets was generated - cells were lysed on ice for 20 minutes with whole cell extract buffer (50 mM HEPES pH 7.4, 150 mM NaCl, 1 mM EDTA, 0.5% NP-40, 10% glycerol) with added inhibitors (PMSF, DTT, Na_3_VO_4_, NaF, Leupeptin, Pepstatin, Aprotinin), before they were spun down at 13,000 rpm for 20 minutes. Supernatant was collected and samples were run on 10% or 15% SDS-PAGE and analyzed by western blot using Clarity Western ECL Blotting Substrates (Bio-Rad Laboratories Inc.) and imaged using a ChemiDoc MP (Bio-Rad Laboratories Inc.). Primary antibodies used: HA (H3663, Sigma, 1:1000), Ku70 (N3H10; Santa Cruz Biotechnology, Inc., 1:1000), Ku80 (M-20; Santa Cruz, 1:500), and mouse α-tubulin (T5168, Sigma, 1:1000). Primary antibodies of proteomic analysis candidates validated by western blot: eIF3η (C-5, Santa Cruz, 1:1000), Myosin VI (A-9, Santa Cruz, 1:100), Pdcd-4 (B-4, Santa Cruz, 1:1000), and Rabbit α-tubulin (ab15246; Abcam, 1:1000). Secondary antibodies were: Peroxidase-conjugated AffiniPure Goat Anti-Mouse IgG (1:5000), mouse anti-goat IgG-HRP (Santa Cruz Biotechnology Inc., 1:3000), goat anti-rabbit IgG (H+L)-HRP Conjugate (BioRad, 1:5000). Western blot samples were quantified using Image Lab 6.0.1. After detection of lanes and bands, adjusted volumes detected for experimental samples were normalized using the Day 1 Dox samples for each blot.

### PCR, T7 Endonuclease Assays, and Sanger Sequencing Validation of Editing

Cells were harvested and DNA was extracted from cells using QuickExtract DNA Extraction Solution (Lucigen Corporation). The pellet was dissolved in 20-80 uL of QuickExtract solution. DNA surrounding target sites 1, 2, and 3 was amplified using PCR (see Supplementary for primers).

T7 Endonuclease I (T7E1) assay was conducted following the extraction of genomic DNA. T7E1 (New England BioLabs Inc.) was used for this assay. PCR amplified DNA from potential knockout clones and wild-type DNA were mixed in a reaction in which DNA was denatured at 95°C for 5 minutes, and then cooled slowly to room temperature to allow DNA from knockout and wild-type samples to anneal together. T7E1 was then added (1 μL) to the annealed PCR products and incubated at 37°C for 15 minutes to allow DNA digestion by the enzyme. T7E1 cuts at mismatches in double-stranded DNA that occur from annealing of edited knockout DNA with wild-type DNA. Restriction enzyme products were visualized via agarose gel electrophoresis to identify evidence of editing in potential Ku70 knockouts.

Following a positive T7 endonuclease assay result, PCR amplified DNA of the target site of interest was purified via a GeneJet PCR Purification kit (ThermoFisher) according to the manufacturer’s instructions. Purified DNA was then sent for Sanger sequencing at the London Regional Genomics Center. SnapGene was used to align CRISPR edited DNA with wild type DNA. DECODR.org was used to validate and assess editing efficiency at target sites.

### Crystal Violet Assays

Control and Ku70 knockout cells were plated onto 96-well plates on Day 5 of Dox treatment or post Dox withdrawal. For each condition 10,000 cells were plated in a 96-well plate (5 wells/replicates per clone and condition). Cells were then fixed in 4% PFA and then incubated in Crystal Violet solution (0.5% in 20% methanol). Pictures were taken, and then 100uL of 2% SDS was added to each well to dissolve the crystal violet dye and plates were left for 30 minutes at room temperature. The BioTek Epoch Microplate Spectrophotometer was used to take readings at 550nm wavelengths using the Gen5 all-in-one platereader program. Graphs were created using GraphPad Prism9. The number of cells adhered to each well was inferred from a standard curve generated by plating known numbers of cells and recording readings at 550nm wavelengths.

Samples and conditions were compared using a two-way ANOVA via Prism9 Matched values stacked in a sub column with interaction term was included. The Geisser-Greenhouse correction was also utilized. Within each row, columns were compared with every other column. Correction for multiple comparisons was done via a Tukey test.

### Immunofluorescence of γH2AX foci

Cells were plated with or without doxycycline depending on the condition. After splitting cells on Days 3 and 5, cells were seeded onto coverslips in a 24-well plate, and returned to the incubator to be fixed on Day 5 and 8 post Dox removal, respectively. Cells were fixed with 4% PFA, and processed for indirect immunofluorescence analysis according to standard protocols using phospho-Histone H2A.X (S139) (20E3) Rabbit antibody (Cell Signaling Technology, 1:1000) and Alexa Fluor 647 goat anti-rabbit IgG (H+L) secondary antibody (Invitrogen, 1:1000). Cells were mounted using ProLong Diamond Antifade Mountant with DAPI (Invitrogen) and imaged the next day using an Olympus BX51 microscope at 40X magnification and Image-Pro Plus software (Media Cybernetics, Inc.). ImageJ was used to quantify the number of γH2AX foci per nucleus. Nuclei were counted manually and denoted by the freehand selections tool and foci were counted using the find maxima tool (brightness for γH2AX foci images set to 0-31, prominence for find maxima set to 2). GaphPad Prism9 was used to generate foci quantification graphs. An ordinary one-way ANOVA with multiple comparisons was used to assess statistical significance of foci quantification. Density plots were created using RStudio.

### Telomere Restriction Fragment (TRF) Analyses

2.2 x 10^6^ cells were plated on 10cm plates for each cell line with DMEM (10% FBS). Cells were plated with or without doxycycline depending on the condition. Cells were split 1:3 on Day 3 and Day 5. Cells were harvested on Day 1 Dox and Day 8 Dox/No Dox. DNA from cell pellets was extracted using the PureLink^TM^ Genomic DNA Mini Kit (Invitrogen) according to the manufacturer’s instructions. Mean telomere lengths of samples were assessed using the TeloTAGGG^TM^ Telomere Length Assay (Roche) according to manufacturer’s instructions aside from modifications listed. For each sample, 4 μg of DNA was digested with Hinf I/Rsa I enzyme mixture. An overnight (14 hour) capillary transfer setup was used to transfer the DNA to BrightStar^TM^ – Plus positively charged nylon membrane (Invitrogen) using 20X SSC transfer buffer. A DNA crosslinker was used to fix the DNA on the nylon membrane following overnight transfer. Chemiluminescent images were taken using a ChemiDoc^TM^ MP Imaging System. ImageLab was used to assess average telomere lengths for each sample. Graphs were created using GraphPad Prism9.

### Proteomic Analysis by Mass Spectrometry

SB Ku70 knockout cells were plated on 6-well plates (300,000 per well) according to depletion curve protocols described above. SB cells were plated in duplicate, with one plate containing +Dox media, and the other -Dox media undergoing Ku70-HA depletion (N=3) Another two plates were also seeded per replicate for western blots. Samples were trypsinized, spun down (8,000 rpm for 3 minutes, washed with PBS, and spun down again) and collected from Day 1 Dox to Day 8 No Dox (or Day 8 Dox for control group). Samples for western blots were prepared as described above, with 35 μg of protein loaded per well to 10% SDS-PAGE gels. Samples for mass spectrometry analysis for global proteomics were prepared exactly as described previously, but following the digestion and acidification, peptides were desalted using Pierce^TM^ C18 Spin Tips (Cat# 84850)[67]. Samples were then dried in a Speed vacuum, resuspended in 0.1% formic acid, and quantified by BCA assay. Approximately 500 ng of peptide sample was injected onto a Waters M-Class nanoAcquity UHPLC system (Waters, Milford, MA) coupled to an ESI Orbitrap mass spectrometer (Q Exactive plus, ThermoFisher Scientific) operated as described in Maitland et al, 2021. All MS raw files were searched in MaxQuant version 1.5.8.3 using the Human Uniprot database (reviewed only; updated July 2020). Missed cleavages were set to 3, cysteine carbamidomethylation (CAM) was set as a fixed modification and oxidation (M), N-terminal acetylation (protein) and deamidation (NQ) were set as variable modifications (max. number of modifications per peptide = 5), and peptide length ≥6. Protein and peptide FDR was left to 0.01 (1%) and decoy database was set to revert. Match between runs was enabled and all other parameters left at default[67]. Protein groups were loaded into Perseus (version 1.6.0.7) and proteins containing peptides only identified by site, matched to reverse, potential contaminant, or had less than 2 unique peptides were removed. After log2 transformation, protein groups were only retained if they had valid values in ≥3 samples in either control or Ku70 knockouts for proteome. For proteome analysis, protein group label-free quantification (LFQ) log2 transformed intensities were used. In all datasets, missing values were imputed using a width of 0.3 and down shift of 1.8, and two-sample t tests were performed in Perseus between control and experimental samples for Days 4, 6, and 7. Proteins were filtered according to day collected, fold-change (≥1.5 FC) and p-value (p≤0.05). These filtered protein lists for Days 4, 6, and 7 (Experimental vs Controls) were used for pathway analysis using Metascape[68]. The mass spectrometry proteomics data have been deposited to the ProteomeXchange Consortium via the PRIDE [1] partner repository with the dataset identifier PXD036297[69]. For review purposes, the data can be accessed using the following Reviewer account details:

Username

Password: uIQq8i20

## Acknowledgements

We would like to thank the Schild-Poulter and Edgell lab members for their feedback, expertise, and support they provided on this project. We would also like to thank the labs of Drs. Murray Junop and David Haniford at Western University for sharing reagents and the help with equipment use.

## Supplementary Information

**S1 Fig.** Targeting Three Exon/Intron Junctions for CRISPR editing. Three gRNAs were designed to target the Exon/Intron junctions of Exons 7, 6, and 12 respectively. SaCas9 or TevCas9 endonucleases were used to induce cleavage at the target sites. Expected cut sites are indicated by red lines in the sequence.

**S2 Fig.** Average γH2AX foci accumulation does not change significantly following Dox withdrawal compared to IR treated cells **A**. The average number of γH2AX foci/nucleus in Ku70 knockout cells on the days indicated in media containing Dox or following Dox withdrawal (no Dox) (N=3). TREx-293 Ku70-HA cells treated with 2 Gy of ionizing radiation (IR) act as a positive control. TREx-293 Ku70-HA cells treated with 2 Gy of ionizing radiation are significantly different from untreated TREx-293 Ku70-HA cells and Ku70 knockout cells as denoted by * symbol, but average foci accumulation in knockout cells is not significantly different from unedited TREx-293 Ku70-HA cells as denoted by ns (Ordinary One-Way ANOVA, multiple comparisons, p<0.001).

**S3 Fig.** Volcano plots of altered proteins following Ku70-HA withdrawal Proteins found to be decreased or increased significantly on **A.** Day 4, **B.** Day 6, and **C.** Day 7 post Dox withdrawal (Fold-change ≥ 1.5; p-value ≤ 0.05). The gray line bisecting the y-axis denotes a p-value of 0.05. Following Ku withdrawal, proteins with a fold-change of 1.5 or greater compared to growth-matched controls for each day are denoted by red points on the volcano plot. Proteins that are decreased compared to controls have a negative log(Fold Change) and proteins that are increased have a positive log(Fold Change).

**S4 Fig.** Western blot validation of proteomic data for EIF3B, MYO6, and PDCD4 candidates in alternate Ku70 knockout clone, Sa11 **A**. Western blot validation of proteomic analysis results for 3 candidate proteins at the days listed post Dox withdrawal for the Sa11 Ku70 knockout clone. All samples are from one experiment, but the top and bottom panels are from 2 different western blots. **B.** Quantification of MYO6, EIF3B, and PDCD4 protein levels relative to alpha-tubulin for (N=3) western blots. +/- indicates the presence or absence of Dox from cell media. Day post Dox withdrawal are indicated.

**S1 Data.** Sanger sequencing data and DECODR analysis confirmation of Ku70 editing at target sites 1 and 2 for Sa11, SB, and TI Ku70 knockout clones

**S2 Data.** List of proteins quantified in at least 3 samples in proteomic analysis by mass spectrometry

**S3 Data.** Lists of proteins found to be significantly changed (≥1.5 fold-change, p-value ≤ 0.05) compared to growth-matched controls on Day 4, Day 6, and Day 7

**S4 Data.** Student’s two-way t-tests for proteomic analysis for Day 4, Day 6, and Day 7

**S5 Data.** Metascape results for protein lists significantly changed (≥1.5 fold-change, p-value ≤ 0.05) compared to growth-matched controls on Day 4, Day 6, and Day 7

## References

1. Lieber MR. The mechanism of DSB repair by the NHEJ. Annu Rev Biochem. 2011;79: 181–211. doi:10.1146/annurev.biochem.052308.093131.The

2. Chang HHY, Pannunzio NR, Adachi N, Lieber MR. Non-homologous DNA end joining and alternative pathways to double-strand break repair. Nat Rev Mol Cell Biol. 2017;18: 495–506. doi:10.1038/nrm.2017.48

3. Mari PO, Florea BI, Persengiev SP, Verkaik NS, Brüggenwirth HT, Modesti M, et al. Dynamic assembly of end-joining complexes requires interaction between Ku70/80 and XRCC4. Proc Natl Acad Sci U S A. 2006;103: 18597–18602. doi:10.1073/PNAS.0609061103/SUPPL_FILE/IMAGE530.GIF

4. Zahid S, El Dahan MS, Iehl F, Fernandez-varela P, Du MH Le, Ropars V, et al. The Multifaceted Roles of Ku70/80. Int J Mol Sci 2021, Vol 22, Page 4134. 2021;22: 4134. doi:10.3390/IJMS22084134

5. Fell VL, Schild-Poulter C. The Ku heterodimer: Function in DNA repair and beyond. Mutat Res - Rev Mutat Res. 2015;763: 15–29. doi:10.1016/j.mrrev.2014.06.002

6. Hsu HL, Yannone SM, Chen DJ. Defining interactions between DNA-PK and ligase IV/XRCC4. DNA Repair (Amst). 2002;1: 225–235. doi:10.1016/S1568-7864(01)00018-0

7. Frit P, Ropars V, Modesti M, Charbonnier JB, Calsou P. Plugged into the Ku-DNA hub: The NHEJ network. Progress in Biophysics and Molecular Biology. Elsevier Ltd; 2019. pp. 62–76. doi:10.1016/j.pbiomolbio.2019.03.001

8. Abbasi S, Parmar G, Kelly RD, Balasuriya N, Schild-Poulter C. The Ku complex: recent advances and emerging roles outside of non-homologous end-joining. Cell Mol Life Sci. 2021; doi:10.1007/s00018-021-03801-1

9. Christie SM, Fijen C, Rothenberg E. V(D)J Recombination: Recent Insights in Formation of the Recombinase Complex and Recruitment of DNA Repair Machinery. Front Cell Dev Biol. 2022;0: 850. doi:10.3389/FCELL.2022.886718

10. Indiviglio SM, Bertuch AA. Ku’s essential role in keeping telomeres intact [Internet]. Proceedings of the National Academy of Sciences of the United States of America. 2009. pp. 12217–12218. doi:10.1073/pnas.0906427106

11. De Lange T. Shelterin-mediated telomere protection. Annu Rev Genet. 2018;52: 223–247. doi:10.1146/annurev-genet-032918-021921

12. Shay JW, Wright WE. Telomeres and telomerase: three decades of progress. Nat Rev Genet. 2019;20: 299–309. doi:10.1038/s41576-019-0099-1

13. Fisher TS, Zakian VA. Ku: A multifunctional protein involved in telomere maintenance. DNA Repair. Elsevier; 2005. pp. 1215–1226. doi:10.1016/j.dnarep.2005.04.021

14. Porter SE, Greenwell PW, Ritchie KB, Petes TD. The DNA-binding protein Hdf1p (a putative Ku homologue) is required for maintaining normal telomere length in Saccharomyces cerevisiae. Nucleic Acids Research. 1996.

15. Melnikova L, Biessmann H, Georgiev P. The Ku protein complex is involved in length regulation of Drosophila telomeres. Genetics. 2005;170: 221–235. doi:10.1534/genetics.104.034538

16. Hsu HL, Gilley D, Galande SA, Prakash Hande M, Allen B, Kim SH, et al. Ku acts in a unique way at the mammalian telomere to prevent end joining. Genes Dev. 2000;14: 2807–2812. doi:10.1101/gad.844000

17. Samper E, Goytisolo FA, Slijepcevic P, Van Buul PPW, Blasco MA. Mammalian Ku86 protein prevents telomeric fusions independently of the length of TTAGGG repeats and the G-strand overhang. EMBO Rep. 2000;1: 244–252. doi:10.1093/embo-reports/kvd051

18. D’Adda di Fagagna F, Hande MP, Tong WM, Roth D, Lansdorp PM, Wang ZQ, et al. Effects of DNA nonhomologous end-joining factors on telomere length and chromosomal stability in mammalian cells. Curr Biol. 2001;11: 1192–1196. doi:10.1016/S0960-9822(01)00328-1

19. Sui J, Zhang S, Chen BPC. DNA-dependent protein kinase in telomere maintenance and protection [Internet]. Cellular and Molecular Biology Letters. BioMed Central Ltd.; 2020. p. 2. doi:10.1186/s11658-020-0199-0

20. Chai W, Ford LP, Lenertz L, Wright WE, Shay JW. Human Ku70/80 associates physically with telomerase through interaction with hTERT. J Biol Chem. 2002;277: 47242–47247. doi:10.1074/jbc.M208542200

21. Li G, Nelsen C, Hendrickson EA. Ku86 is essential in human somatic cells. Proc Natl Acad Sci U S A. 2002;99: 832–837. doi:10.1073/pnas.022649699

22. Wang Y, Ghosh G, Hendrickson EA. Ku86 represses lethal telomere deletion events in human somatic cells. Proc Natl Acad Sci U S A. 2009;106: 12430– 12435. doi:10.1073/pnas.0903362106

23. Mazzucco G, Huda A, Galli M, Piccini D, Giannattasio M, Pessina F, et al. Telomere damage induces internal loops that generate telomeric circles. Nat Commun. 2020;11. doi:10.1038/s41467-020-19139-4

24. Nussenzweig A, Chen C, Soares VDC, Sanchez M, Sokol K, Nussenzweig MC, et al. Requirement for Ku80 in growth and immunoglobulin V(D)J recombination. Nature. 1996;382: 551–555. doi:10.1038/382551a0

25. Gu Y, Seidl KJ, Rathbun GA, Zhu C, Manis JP, Van Der Stoep N, et al. Growth retardation and leaky SCID phenotype of Ku70-deficient mice. Immunity. 1997;7: 653–665. doi:10.1016/S1074-7613(00)80386-6

26. De Sena-Tomas C, Yu EY, Calzada A, Holloman WK, Lue NF, Perez-Martin J. Fungal Ku prevents permanent cell cycle arrest by suppressing DNA damage signaling at telomeres. Nucleic Acids Res. 2015;43: 2138–2151. doi:10.1093/nar/gkv082

27. Ghosh G, Li G, Myung K, Hendrickson EA. The Lethality of Ku86 (XRCC5) Loss-of-Function Mutations in Human Cells is Independent of p53 (TP53). Radiat Res. 2007;167: 66–79. doi:10.1667/rr0692.1

28. Uegaki K, Adachi N, So S, Iiizumi S, Koyama H. Heterozygous inactivation of human Ku70/Ku86 heterodimer does not affect cell growth, double-strand break repair, or genome integrity. DNA Repair (Amst). 2006;5: 303–311. doi:10.1016/j.dnarep.2005.10.008

29. Fattah KR, Ruis BL, Hendrickson EA. Mutations to Ku reveal differences in human somatic cell lines. 2008; doi:10.1016/j.dnarep.2008.02.008

30. Saydam O, Saydam N. Deficiency of Ku Induces Host Cell Exploitation in Human Cancer Cells. Front Cell Dev Biol. 2021;9: 1–9. doi:10.3389/fcell.2021.651818

31. Wolfs JM, Hamilton TA, Lant JT, Laforet M, Zhang J, Salemi LM, et al. Biasing genome-editing events toward precise length deletions with an RNA-guided TevCas9 dual nuclease. Proc Natl Acad Sci U S A. 2016;113: 14988–14993. doi:10.1073/pnas.1616343114

32. Myung K, Ghosh G, Fattah FJ, Li G, Kim H, Dutia A, et al. Regulation of Telomere Length and Suppression of Genomic Instability in Human Somatic Cells by Ku86. Mol Cell Biol. 2004;24: 5050–5059. doi:10.1128/mcb.24.11.5050-5059.2004

33. Jaco I, Muñoz P, Blasco MA. Role of human Ku86 in telomere length maintenance and telomere capping. Cancer Res. 2004;64: 7271–7278. doi:10.1158/0008-5472.CAN-04-1381

34. Wang Y, Ghosh G, Hendrickson EA. Ku86 represses lethal telomere deletion events in human somatic cells. Proc Natl Acad Sci U S A. 2009;106: 12430– 12435. doi:10.1073/pnas.0903362106

35. Pansare K, Raj Singh S, Chakravarthy V, Gupta N, Hole A, Gera P, et al. Raman Spectroscopy: An Exploratory Study to Identify Post-Radiation Cell Survival. https://doi.org/101177/0003702820908352. 2020;74: 553–562. doi:10.1177/0003702820908352

36. Slattery SS, Wang H, Giguere DJ, Kocsis C, Urquhart BL, Karas BJ, et al. Plasmid-based complementation of large deletions in Phaeodactylum tricornutum biosynthetic genes generated by Cas9 editing. Sci Reports 2020 101. 2020;10: 1–12. doi:10.1038/s41598-020-70769-6

37. Wang Y, Ghosh G, Hendrickson EA. Ku86 represses lethal telomere deletion events in human somatic cells [Internet]. Proceedings of the National Academy of Sciences of the United States of America. 2009. doi:10.1073/pnas.0903362106

38. Rossiello F, Jurk D, Passos JF, d’Adda di Fagagna F. Telomere dysfunction in ageing and age-related diseases. Nat Cell Biol 2022 242. 2022;24: 135–147. doi:10.1038/s41556-022-00842-x

39. Li B, Reddy S, Comai L. Depletion of Ku70/80 reduces the levels of extrachromosomal telomeric circles and inhibits proliferation of ALT cells. Aging (Albany NY). 2011;3: 395. doi:10.18632/AGING.100308

40. He B, Lu N, Zhou Z. Cellular and Nuclear Degradation during Apoptosis. Curr Opin Cell Biol. 2009;21: 900. doi:10.1016/J.CEB.2009.08.008

41. Xu X, Shi Y chang, Gao W, Mao G, Zhao G, Agrawal S, et al. The novel presenilin-1-associated protein is a proapoptotic mitochondrial protein. J Biol Chem. 2002;277: 48913–48922. doi:10.1074/jbc.M209613200

42. Lamarca V, Sanz-Clemente A, Pérez-Pé R, Martínez-Lorenzo MJ, Halaihel N, Muniesa P, et al. Two isoforms of PSAP/MTCH1 share two proapoptotic domains and multiple internal signals for import into the mitochondrial outer membrane. Am J Physiol - Cell Physiol. 2007;293. doi:10.1152/AJPCELL.00431.2006/ASSET/IMAGES/LARGE/ZH00100753840009.JPEG

43. Bensaad K, Tsuruta A, Selak MA, Vidal MNC, Nakano K, Bartrons R, et al. TIGAR, a p53-inducible regulator of glycolysis and apoptosis. Cell. 2006;126: 107–120. doi:10.1016/J.CELL.2006.05.036

44. Lee P, Vousden KH, Cheung EC. TIGAR, TIGAR, burning bright. Cancer Metab 2014 21. 2014;2: 1–9. doi:10.1186/2049-3002-2-1

45. Widden H, Placzek WJ. The multiple mechanisms of MCL1 in the regulation of cell fate. Commun Biol 2021 41. 2021;4: 1–12. doi:10.1038/s42003-021-02564-6

46. Das S, Maitra U. Functional significance and mechanism of eIF5-promoted GTP hydrolysis in eukaryotic translation initiation. Prog Nucleic Acid Res Mol Biol. 2001;70: 207–231. doi:10.1016/S0079-6603(01)70018-9

47. Ma F, Li X, Ren J, Guo R, Li Y, Liu J, et al. Downregulation of eukaryotic translation initiation factor 3b inhibited proliferation and metastasis of gastric cancer. Cell Death Dis 2019 109. 2019;10: 1–12. doi:10.1038/s41419-019-1846-0

48. Song S, Liu J, Zhang M, Gao X, Sun W, Liu P, et al. Eukaryotic translation initiation factor 3 subunit B could serve as a potential prognostic predictor for breast cancer. Bioengineered. 2022;13: 2762–2776. doi:10.1080/21655979.2021.2017567/SUPPL_FILE/KBIE_A_2017567_SM8035.ZIP

49. Miyoshi M, Okajima T, Matsuda T, Fukuda MN, Nadano D. Bystin in human cancer cells: intracellular localization and function in ribosome biogenesis. Biochem J. 2007;404: 373. doi:10.1042/BJ20061597

50. Young CL, Karbstein K. The roles of S1 RNA-binding domains in Rrp5’s interactions with pre-rRNA. RNA. 2011;17: 512. doi:10.1261/RNA.2458811

51. Piñeiro D, Stoneley M, Ramakrishna M, Alexandrova J, Dezi V, Juke-Jones R, et al. Identification of the RNA polymerase I-RNA interactome. Nucleic Acids Res. 2018;46: 11002–11013. doi:10.1093/NAR/GKY779

52. Shao Z, Flynn RA, Crowe JL, Zhu Y, Liang J, Jiang W, et al. DNA-PKcs has KU-dependent function in rRNA processing and haematopoiesis. Nature. 2020;579: 291–296. doi:10.1038/s41586-020-2041-2

53. Nelson HA, Roe MW. Molecular physiology and pathophysiology of stromal interactionmolecules. Exp Biol Med. 2018;243: 451. doi:10.1177/1535370218754524

54. Ibsen MS, Gad HH, Thavachelvam K, Boesen T, Desprès P, Hartmann R. The 2′-5′-Oligoadenylate Synthetase 3 Enzyme Potently Synthesizes the 2′-5′-Oligoadenylates Required for RNase L Activation. J Virol. 2014;88: 14222. doi:10.1128/JVI.01763-14

55. Anisenko AN, Knyazhanskaya ES, Zatsepin TS, Gottikh MB. Human Ku70 protein binds hairpin RNA and double stranded DNA through two different sites. Biochimie. 2017;132: 85–93. doi:10.1016/J.BIOCHI.2016.11.001

56. Shiba Y, Randazzo PA. ArfGAP1 function in COPI mediated membrane traffic: Currently debated models and comparison to other coat-binding ArfGAPs. Histol Histopathol. 2012;27: 1143. doi:10.14670/HH-27.1143

57. Li J, D’Angiolella V, Seeley ES, Kim S, Kobayashi T, Fu W, et al. USP33 regulates centrosome biogenesis via de-ubiquitylation of a centriolar protein, CP110. Nature. 2013;495: 255–259. doi:10.1038/NATURE11941

58. Muñoz P, Zdzienicka MZ, Blanchard J-M, Piette J. Hypersensitivity of Ku-Deficient Cells toward the DNATopoisomerase II Inhibitor ICRF-193 Suggests a Novel Role for KuAntigen during the G2 and M Phases of the CellCycle. Mol Cell Biol. 1998;18: 5797. doi:10.1128/MCB.18.10.5797

59. Rampakakis E, Di Paola D, Zannis-Hadjopoulos M. Ku is involved in cell growth, DNA replication and G1-S transition. J Cell Sci. 2008;121: 590–600. doi:10.1242/jcs.021352

60. You W, Tan G, Sheng N, Gong J, Yan J, Chen D, et al. Downregulation of myosin VI reduced cell growth and increased apoptosis in human colorectal cancer. Acta Biochim Biophys Sin (Shanghai). 2016;48: 430. doi:10.1093/ABBS/GMW020

61. Wang D, Zhu L, Liao M, Zeng T, Zhuo W, Yang S, et al. MYO6 knockdown inhibits the growth and induces the apoptosis of prostate cancer cells by decreasing the phosphorylation of ERK1/2 and PRAS40. Oncol Rep. 2016;36: 1285–1292. doi:10.3892/OR.2016.4910/HTML

62. O’Loughlin T, Masters TA, Buss F. The MYO6 interactome reveals adaptor complexes coordinating early endosome and cytoskeletal dynamics. EMBO Rep. 2018;19: e44884. doi:10.15252/EMBR.201744884

63. Eto K, Goto S, Nakashima W, Ura Y, Abe SI. Loss of programmed cell death 4 induces apoptosis by promoting the translation of procaspase-3 mRNA. Cell Death Differ 2012 194. 2011;19: 573–581. doi:10.1038/cdd.2011.126

64. Ran FA, Hsu PD, Wright J, Agarwala V, Scott DA, Zhang F. Genome engineering using the CRISPR-Cas9 system. Nat Protoc 2013 811. 2013;8: 2281–2308. doi:10.1038/nprot.2013.143

65. Strathdee CA, McLeod MR, Hall JR. Efficient control of tetracycline-responsive gene expression from an autoregulated bi-directional expression vector. Gene. 1999;229: 21–29. doi:10.1016/S0378-1119(99)00045-1

66. Engler C, Kandzia R, Marillonnet S. A One Pot, One Step, Precision Cloning Method with High Throughput Capability. PLoS One. 2008;3: e3647. doi:10.1371/JOURNAL.PONE.0003647

67. Maitland MER, Kuljanin M, Wang X, Lajoie GA, Schild-Poulter C. Proteomic analysis of ubiquitination substrates reveals a CTLH E3 ligase complex-dependent regulation of glycolysis. FASEB J. 2021;35. doi:10.1096/FJ.202100664R

68. Zhou Y, Zhou B, Pache L, Chang M, Khodabakhshi AH, Tanaseichuk O, et al. Metascape provides a biologist-oriented resource for the analysis of systems-level datasets. Nat Commun. 2019;10. doi:10.1038/S41467-019-09234-6

69. Perez-Riverol Y, Csordas A, Bai J, Bernal-Llinares M, Hewapathirana S, Kundu DJ, et al. The PRIDE database and related tools and resources in 2019: Improving support for quantification data. Nucleic Acids Res. 2019;47: D442– D450. doi:10.1093/nar/gky1106

